# Analysis of allelic series with transcriptomic phenotypes

**DOI:** 10.1101/210724

**Authors:** David Angeles-Albores, Paul W. Sternberg

## Abstract

Although transcriptomes have recently been used to perform epistasis analyses, they are not yet used to study intragenic function/structure relationships. We developed a theoretical framework to study allelic series using transcriptomic phenotypes. As a proof-of-concept, we apply our methods to an allelic series of *dpy-22*, a highly pleiotropic *Caenorhabditis elegans* gene orthologous to the human gene *MED12*, which is a subunit of the Mediator complex. Our methods identify functional regions within *dpy-22* that modulate Mediator activity upon various genetic modules.

## Introduction

Mutations of a gene can yield a series of alleles with different phenotypes that reveal multiple functions encoded by that gene, regardless of the alleles’ molecular nature. Homozygous alleles can be ordered by their phenotypic severity; then, phenotypes of *trans-*heterozygotes carrying two alleles can reveal which alleles are dominant for each phenotype. Together, the severity and dominance hierarchies show intra-genic functional regions. In *Caenorhabditis elegans*, these series have helped characterize genes such as *let-23/EGFR, lin-3/EGF* and *lin-12/NOTCH* ^1,2,3^

Biology has moved from expression measurements of single genes towards genome-wide measurements. Expression profiling via RNA-seq^4^ enables simultaneous measurement of transcript levels for all genes in a genome, yielding a transcriptome. These measurements can be made on whole organisms, isolated tissues, or single cells^5,6^. Transcriptomes have been successfully used to identify new cell or organismal states^7,8^. For mutant genes, transcriptomic states can be used for epistasis analysis^9,10^, but have not been used to characterize allelic series.

We have devised methods for characterizing allelic series with RNA-seq. To test these methods, we selected three alleles^11,12^ of a *C. elegans* Mediator complex subunit gene, *dpy-22*. Mediator is a macromolecular complex with ∼ 25 subunits^13^ that globally regulates RNA polymerase II (Pol II)^14,15^. The Mediator complex has at least four biochemically distinct modules: the Head, Middle and Tail modules and a CDK-8-associated Kinase Module (CKM).

The CKM associates reversibly with other modules, and appears to inhibit transcription^16,17^. In *C. elegans* development, the CKM promotes both male tail formation^11^ (through interactions with the Wnt pathway), and vulval formation^18^ (through inhibition of the Ras pathway). Homozygotes of allele *dpy-22(bx93)*, which encodes a premature stop codon Q2549Amber^11^, appear grossly wild-type. In contrast, animals homozygous for a more severe allele, *dpy-22(sy622)* encoding another premature stop codon, Q1698Amber^12^, are dumpy (Dpy), have egg-laying defects (Egl), and have multiple vulvae (Muv). (see Fig. 1A). In spite of its causative role in a number of neurodevelopmental disorders^19^, the structural and functional features of this gene are poorly understood. In humans, MED12 is known to have a proline-, glutamine-and leucine-rich domain that interacts with the WNT pathway^20^. However, many disease-causing variants fall outside of this domain^21^. To study these variants and how they interfere with the functionality of *MED12*, quantitative and efficient methods are necessary.

RNA-seq phenotypes have the potential to reveal functional regions within genes, but their phenotypic complexity makes this difficult. We developed a method for determining allelic series from transcriptomic phenotypes and used the *C. elegans dpy-22* gene as a test case. Our analysis revealed functional regions that act to modulate Mediator activity at thousands of genetic loci.

## Results and Discussion

We adapted the allelic series method, previously used for individual phenotypes, for use with expression profiles as multidimensional phenotypes (see Fig. 1). As a proof of principle, we carried out RNA-seq on biological triplicates of mRNA extracted from *dpy-22(sy622)* homozygotes, *dpy-22(bx93)* homozygotes and wild type controls, along with quadruplicates from *trans*-heterozygotes of both alleles. Sequencing was performed at a depth of 20 million reads per sample. Reads were pseudoaligned using Kallisto^22^. We performed a differential expression using a general linear model specified using Sleuth^23^ (see Methods). Differential expression with respect to the wild type control for each transcript *i* in a genotype *g* is measured via a coefficient *ß_g,i_*, which can be loosely interpreted as the natural logarithm of the fold-change. Transcripts were considered to have differential expression between wild-type and a mutant if the false discovery rate, *q*, was less than or equal to 10%. Supplementary File 1 contains all the beta values associated with this project. We have also generated a website containing complete details of all the analyses available at the following URL: https://wormlabcaltech.github.io/med-cafe/analysis.

By these criteria, we found 481 genes differentially expressed in *dpy-22(bx93)* homozygotes, and 2,863 differentially expressed genes in *dpy-22(sy622)* homozygotes (see Basic Statistics Notebook). *Trans*-heterozygotes with the genotype *dpy-6(e14) dpy-22(bx93)/+ dpy-22(sy622)* had 2,214 differentially expressed genes with respect to the wild type.

We used a false hit analysis to identify four non-overlapping phenotypic classes. We use the term genotype-specific to refer to groups of transcripts that were perturbed in one mutant. We use the term genotype-associated to refer to those groups of transcripts whose expression was significantly altered in two or more mutants with respect to the wild type control. The ***dpy-22(sy622)-associated*** phenotypic class consisted of 720 genes differentially expressed in *dpy-22(sy622)* homozygotes and in *trans-*heterozygotes, but which had wild-type expression in *dpy-22(bx93)* homozygotes. The ***dpy-22(bx93)-***associated phenotypic class contains 403 genes differentially expressed in all genotypes. We also identified a ***dpy-22(sy622)-specific*** phenotypic class (1,841 genes) and a trans-heterozygote-specific phenotypic class (1,226 genes; see the Phenotypic Classes Notebook). All genotype-associated phenotypes had Spearman rank correlations > 0.8, indicating that transcripts within these classes changed in the same direction amongst the genotypes studied.

**Table 1.** Dominance analysis for the *dpy-22/MDT12* allelic series. Dominance values closer to 1 indicate *dpy-22(bx93)* is dominant over *dpy-22(sy622)*, whereas 0 indicates *dpy-22(sy622)* is dominant over *dpy-22(bx93).*

| Phenotypic Class | Dominance |
| --- | --- |
| *dpy-22(sy622)*-specific | $1.00 \pm 0.00$ |
| *dpy-22(sy622)*-associated | $0.51 \pm 0.01$ |
| *dpy-22(bx93)*-associated | $0.81 \pm 0.01$ |

We measured allelic dominance for each class using a dominance coefficient (see Methods). The dominance coefficient is a measure of the contribution of each allele to the total expression level in *trans-*heterozygotes. By definition, the *dpy-22(sy622)* allele is completely recessive to *dpy-22(bx93)* for the *dpy-22(sy622)-speciñc* phenotypic class. The *dpy-22(sy622)* and *dpy-22(bx93)* alleles are semidominant *(d_bx93_* = 0.51) to each other for the *dpy-22(sy622*)-associated phenotypic class. The *dpy-22(bx93)* allele is largely dominant over the *dpy-22(sy622)* allele *(d_bx93_* = 0.81; see Table 1) for the *dpy-22(bx93*)-associated phenotypic class.

Because the mutations we used are truncations, our results suggest the existence of various functional regions in *dpy-22/MDT12* (see Fig. 2). The *dpy-22(sy622)-specific* phenotypic class is likely controlled by a single functional region, functional region 1 (FC1), and the *dpy-22(sy622*)-associated phenotypic class is likely controlled by a second functional region, functional region 2 (FC2). It is unlikely that these regions are identical because their dominance behaviors are very different. The *dpy-22(bx93)* allele was largely dominant over the *dpy-22(sy622)* allele for the *dpy-22(bx93*)-associated class, but gene expression in this class was perturbed in both homozygotes. The perturbations were greater for *dpy-22(sy622)* homozygotes than for *dpy-22(bx93)* homozygotes. This behavior can be explained if the *dpy-22(bx93)-*associated class is controlled jointly by two distinct effectors, functional regions 3 and 4 (FC3, FR4, see Fig. 2). A rigorous examination of this model will require studying alleles that mutate the region between Q1689 and Q2549 using homozygotes and *trans*-heterozygotes.

We also found a class of transcripts that had perturbed levels in *trans*-heterozygotes only; its biological significance is unclear. Phenotypes unique to *trans*-heterozygotes are often the result of physical interactions such as homodimerization, or dosage reduction of a toxic product^24^. In the case of *dpy-22/MDT12* orthologs, how either mechanism could operate is not obvious, since DPY-22 is expected to assemble in a monomeric manner into the CKM. Massive single-cell RNA-seq of *C. elegans* has recently been reported^25^. When this technique becomes cost-efficient, single-cell profiling of these genotypes may provide information that complements the whole-organism expression phenotypes, perhaps explaining the origin of this phenotype.

**Figure 1.**
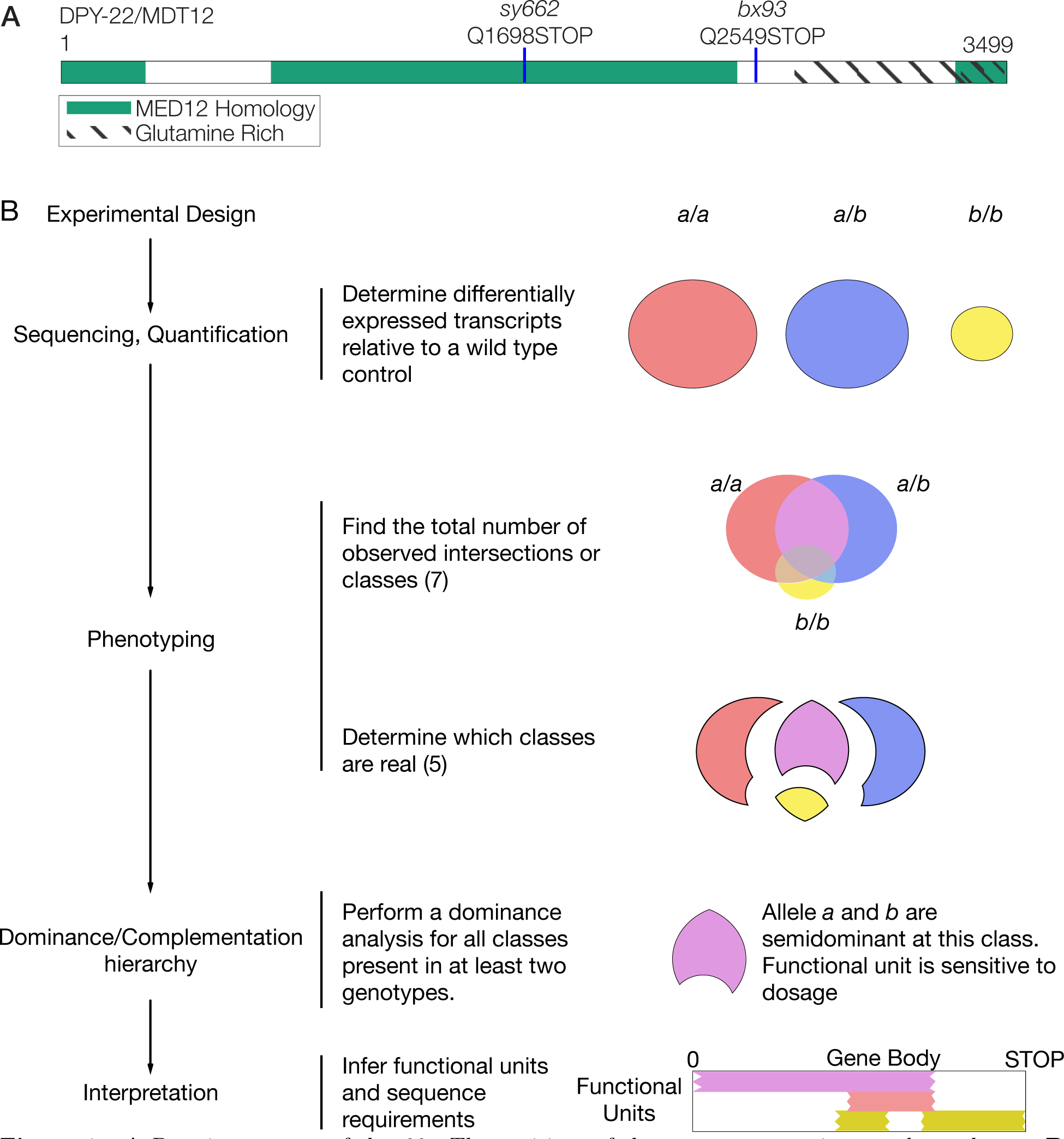
A Protein sequence of *dpy-22.* The positions of the nonsense mutations used are shown. B Flowchart for an analysis of arbitrary allelic series. A set of alleles is selected, and the corresponding genotypes are sequenced. Independent phenotypic classes are then identified. For each phenotypic class, the alleles are ordered in a dominance/complementation hierarchy, which can then be used to infer functional regions within the genes in question.

**Figure 2.**
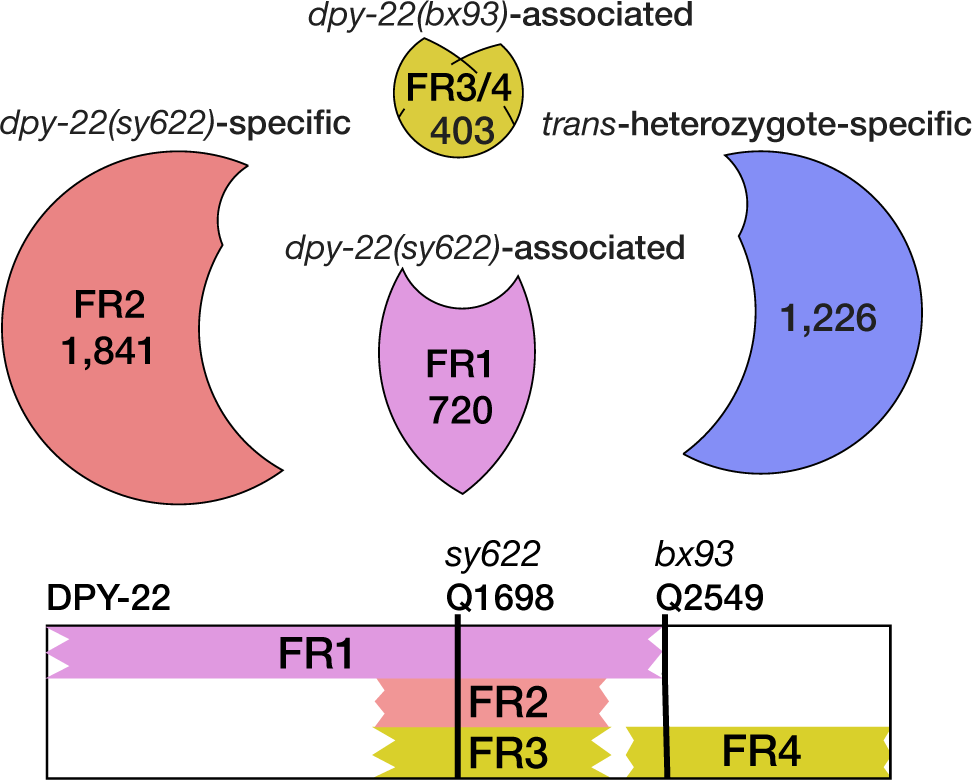
The functional regions associated with each phenotypic class can be mapped intragenically. The number of genes associated with each class is shown. The *dpy-22(bx93*)-associated class may be controlled by two functional regions. FR2 and FR3 could be redundant if FR4 is a modifier of FR2 functionality at *dpy-22(bx93*)-associated loci. Note that the *dpy-22(bx93*)-associated phenotypic class is actually three classes merged together. Two of these classes are DE in *dpy-22(bx93)* homozygotes and one other genotype. Our analyses suggested that these two classes are likely the result of false negative hits and genes in these classes should be differentially expressed in all three genotypes, so they we merged all classes together (see Methods).

Intragenic mapping of functional regions associated with phenotypic classes is important, but their biological meaning remains unclear. To assign biological functionality to phenotypic classes, we extracted transcriptomic signatures associated with a Dumpy (Dpy) phenotype using transcriptomes from *dpy-7* and *dpy-10* mutants (DAA, CPR and PWS *unpublished)*, and a *hif-1*-dependent hypoxia response from a previously published analysis^10^ and asked whether any phenotypic class was enriched in either response. The *sy622*-specific and-associated classes were enriched in genes that are transcriptionally associated with a Dpy phenotype (fold-change enrichment = 3, *p* = 2 ⋅ 1010^−40^, 167 genes observed; fold-change = 1.9, *p* = 9⋅1010^−9^, 82 genes observed). The *bx93*-associated class also showed significant enrichment (fold-change = 2.2, *p* = 4 ⋅ 1010^−10^, 68 genes observed). The class that showed the most extreme deviation from random was the *sy622*-specific class. *dpy-22(sy622)* homozygotes are severely Dpy, whereas *dpy-22(bx93)* homozygotes and *trans*-heterozygotes have a slight Dpy phenotype. Plotting the changes in gene expression for *sy622* homozygotes versus the changes in expression in *dpy-7* mutants revealed that 75% of the transcripts were strongly correlated in both genotypes (see Figure 3). Therefore, the *sy622*-specific phenotypic class contains a transcriptional signature associated with morphological Dpy phenotype (see the Enrichment Notebook).

*dpy-22* is not known to be upstream of the *hif-1*-dependent hypoxia response in *C. elegans.* Enrichment tests revealed that the hypoxia response was significantly enriched in the *bx93*-associated (fold-change = 2.1, *p* = 10^−8^, 63 genes observed), the *sy622*-associated (fold-change = 1.9, *p* = 4 ⋅ 1010^−8^, 78 genes observed) and the *sy622*-specific classes (fold-change = 2.4, *p* = 9 ⋅ 1010^−55^, 186 genes observed). However, there was no correlation between the expression levels of these genes in *dpy-22* genotypes and the expression levels expected from the hypoxia response. Although the hypoxia gene battery can be found in *dpy-22* mutants, these genes are not used to deploy a *hif-1*-dependent hypoxia phenotype. Taken together, our results suggest that transcriptomic signatures can be used to understand the biological functionality of phenotypic classes, and they may be useful in associating phenotypic classes with other phenotypes. This highlights the importance of generating an index set of mutants that can be used to derive a gold standard of transcriptional signatures with which to test future results.

Transcriptomic phenotypes generate large amounts of differential gene expression data, so false positive and false negative rates can lead to spurious phenotypic classes whose putative biological significance is badly misleading. Such artifacts are particularly likely for small phenotypic classes, which should be viewed with skepticism. Notably, errors of interpretation cannot be avoided by setting a more stringent *q-*value cut-off: doing so will decrease the false positive rate, but increase the false negative rate, which will in turn produce smaller phenotypic classes than expected. Our method avoids this pitfall by using total error rate estimates to assess the plausibility of each class. These conclusions are of broad significance to research where highly multiplexed measurements are compared to identify similarities and differences in the genome-wide behavior of a single variable under multiple conditions.

We have shown that transcriptomes can be used to study allelic series in the context of a large, pleiotropic gene. We identified separable phenotypic classes that would otherwise be obscured by other methods, correlated each class to a functional region, and identified sequence requirements for each region. Given the importance of allelic series for characterizing gene function and their roles in specific genetic pathways, we are optimistic that this method will be a useful addition to the geneticist’s arsenal.

## Methods

### Strains used

Strains used were N2 wild-type (Bristol), PS4087 *dpy-22(sy622)*, PS4187 *dpy-22(bx93)*, and PS4176 *dpy-6(e14) dpy-22(bx93)/ + dpy-22(sy622).* Lines were grown on standard nematode growth media (NGM) Petri plates seeded with OP50 *E. coli* at 20°C^26^.

### Strain synchronization, harvesting and RNA sequencing

Strains were synchronized by bleaching Po’s into virgin S. basal (no cholesterol or ethanol added) for 8-12 hours. Arrested L1 larvae were placed in NGM plates seeded with OP50 at 20°C and grown to the young adult stage (assessed by vulval morphology and lack of embryos). RNA extraction and sequencing was performed as previously described by Angeles-Albores *et al*^10,7^.

**Figure 3.**
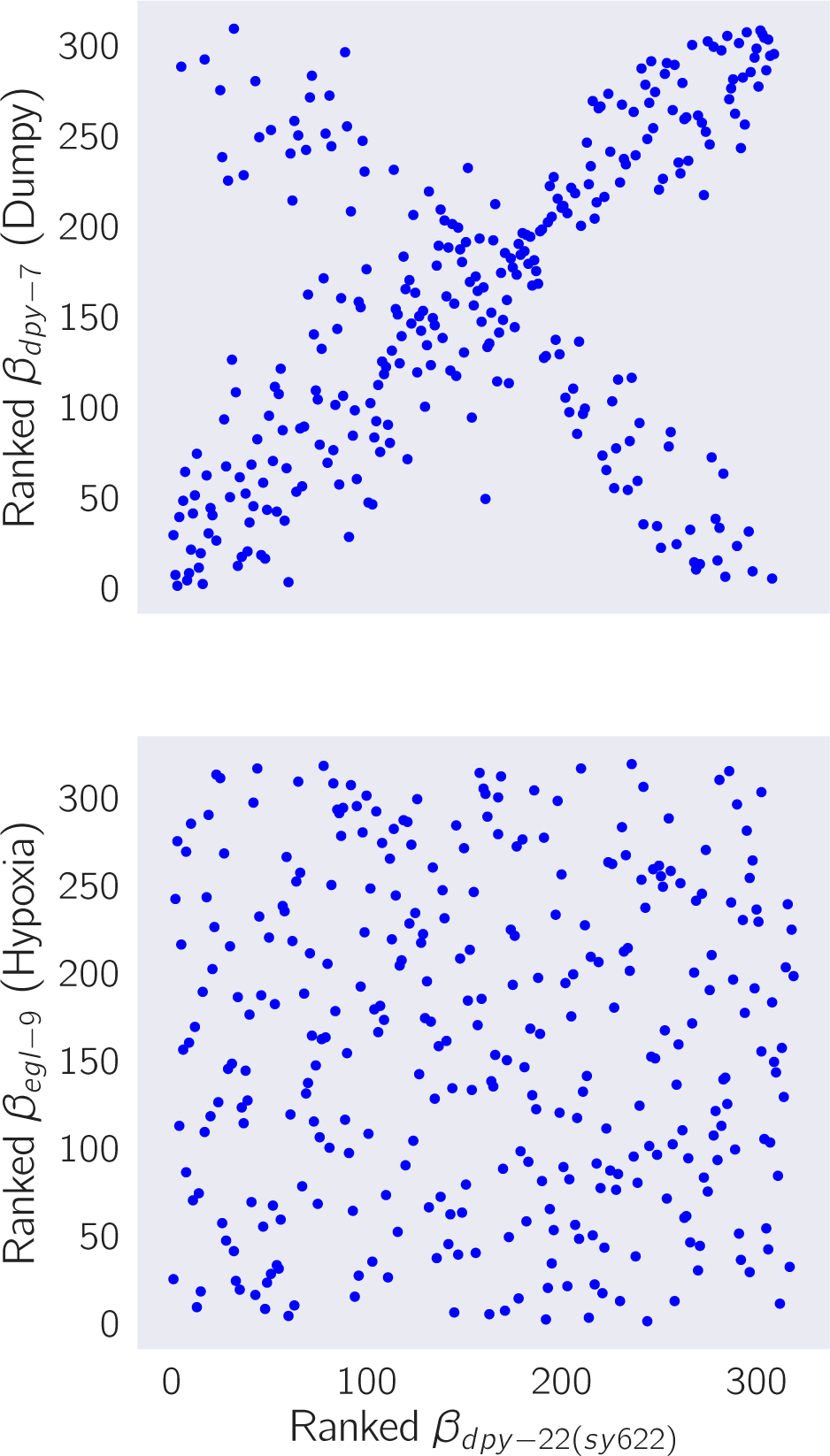
*sy622* homozygotes show a transcriptional response associated with the Dpy phenotype. A We obtained a set of transcripts associated with the Dpy phenotype from *dpy-7* and *dpy-10* mutants. We identified the transcripts that were differentially expressed in *sy622* homozygotes. We ranked the *β* values of each transcript in *sy622* homozygotes and plotted them against the ranked *β* values in *dpy-7* mutants. A significant portion of the genes are correlated between the two genotypes, showing that the signature is largely intact. 25% of the genes are anti-correlated. B We performed the same analysis using a set of transcripts associated with the *hif-1-*dependent hypoxia response as a negative control. Although *sy622* is enriched for the transcripts that make up this response, there is no correlation between the *β* values in *sy622* homozygotes and the *β* values in *egl-9* homozygotes.

### Read pseudo-alignment and differential expression

Reads were pseudo-aligned to the *C. elegans* genome (WBcel235) using Kallisto^22^, using 200 bootstraps and with the sequence bias (--`seqBias`) flag. The fragment size for all libraries was set to 200 and the standard deviation to 40. Quality control was performed on a subset of the reads using FastQC, RNAseQC, BowTie and MultiQC^27,28,29,30^.

Differential expression analysis was performed using Sleuth^23^. We used a general linear model to identify genes that were differentially expressed between wild-type and mutant libraries. To increase our statistical power, we pooled young adult wild-type replicates from other published^10,7^ and unpublished analyses adjusting for batch effects.

### False hit analysis

To accurately count phenotypes, we developed a false hit algorithm (Algorithm 1). We implemented this algorithm for three-way comparisons in Python. Although experimentally restricted, a three-way comparison can result in > 5, 000 possible sets (ignoring size). This large number of models necessitates an algorithmic approach that can at least restrict the possible number of models. Our algorithm uses a noise function that assumes false hit events are non-overlapping (i.e. the same gene cannot be the result of two false positive events in two or more genotypes) to determine the average noise flux between phenotypic classes. These assumptions break down rapidly if false-positive or negative rates exceed 20%.

To benchmark our algorithm, we generated one thousand Venn diagrams at random. For each Venn diagram, we calculated the average false positive and false negative flux matrices. Then, we added noise to each phenotypic class in the Venn diagram, assuming that fluxes were normally distributed with mean and standard deviation equal to the flux coefficient calculated. We input the noised Venn diagram into our false hit analysis and collected classification statistics. For a given signal-to-noise cutoff, λ, classification accuracy varied significantly with changes in the total error rate. In the absence of false negative hits, false hit analysis can accurately identify non-empty genotype-associated phenotypic classes, but identifying genotype-specific classes becomes difficult if the experimental false positive rate is high. On the other hand, even moderate false negative rates (> 10%) rapidly degrade signal from genotype-associated classes. For classes that are associated with three genotypes, an experimental false negative rate of 30% is enough on average to prevents this class from being observed.

We selected λ = 3 because classification using this threshold was high across a range of false positive and false negative combinations. A challenge to applying this algorithm to our data is the fact that the false negative rate for our experiment is unknown. Although there has been significant progress in controlling and estimating false positive rates, we know of no such attempts for false negative rates. It is unlikely that the false negative rate for our study is lower than the false positive rate, because all genotypes except the controls are likely underpowered. We used false negative rates between 10-20% for false hit analysis. When the false negative rate was set at 15% or higher, the algorithm converged on the same five classes shown above. For false negative rates between 10-15%, the algorithm output the same five classes, but also accepted the (*dpy-22(sy622),dpy-22(bx93)*)-associated class. We selected the model corresponding to false negative rates of 15-20% because this model had lower *χ*^2^ values than the model selected with a false negative rate of 10-15% (4,212 versus 100,650).

We asked whether re-classification of some classes into others could improve model fit. We manually re-classified the *(dpy-22(sy622),dpy-22(bx93))-*associated and the *(dpy-22(bx93), trans-heterozygote*)-associated classes into the *bx93-*associated class (which is associated with all genotypes), and we compared *χ*^2^ statistics between a re-classified reduced model and a reduced model. The re-classified model had a lower *χ*^2^ (181). Thus, we concluded that the re-classified reduced model is the most likely model to give rise to our data.

**Data**: **M**_*obs*_ = {*N_l_*}, an observed set of classes, where each class is labelled by *l* ∈ *L* and is of size *N_l_*. *f_p_*, *f_n_*, the false positive and negative rates respectively. *α*, the signal-to-noise threshold for acceptance of a class.

**Result**: **M***_reduced_*, a reduced model that fits the data.

**Algorithm 1:**
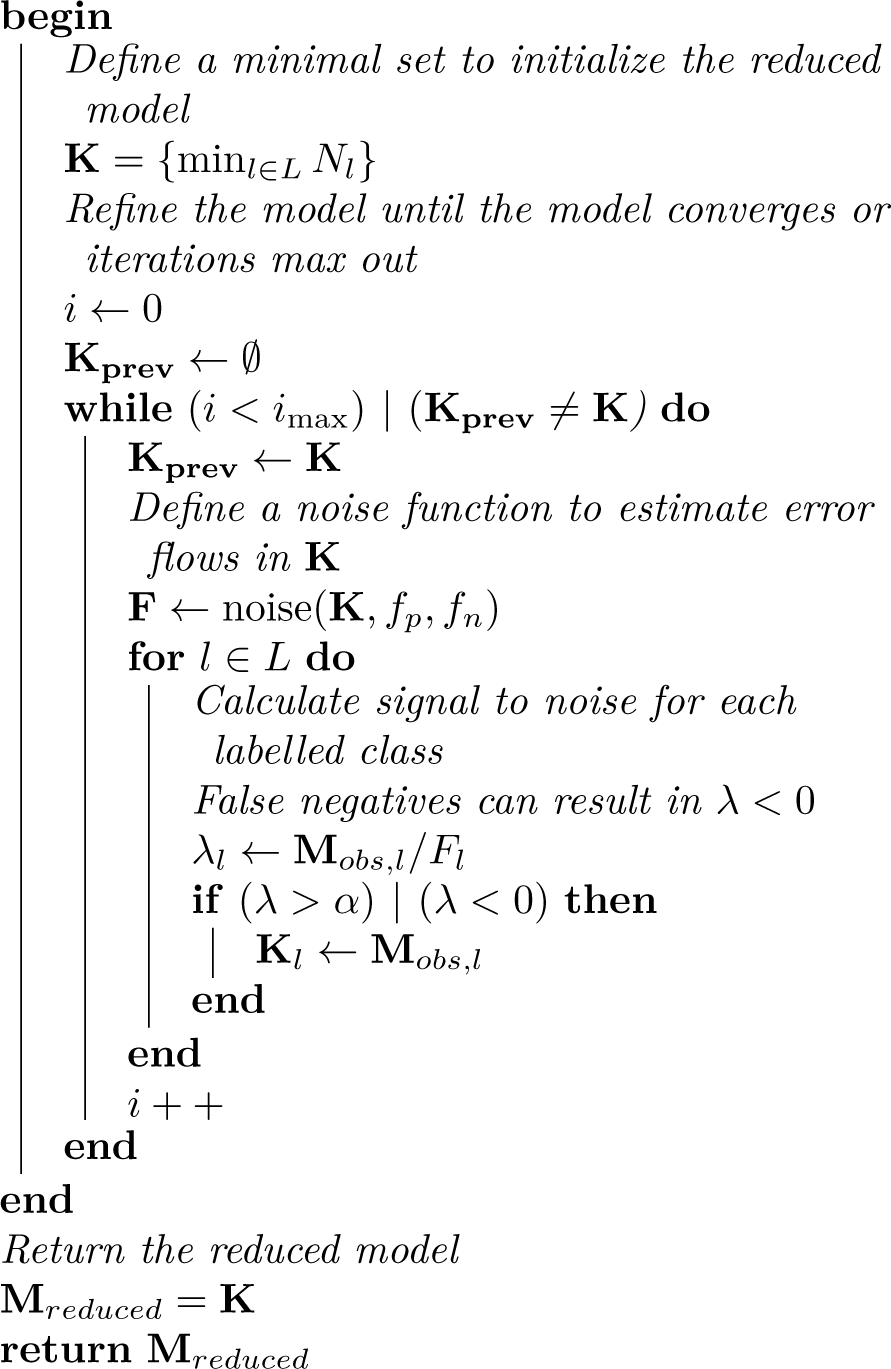
False Hit Algorithm. Briefly, the algorithm initializes a reduced model with the phenotypic class or classes labelled by the largest number of genotypes. This reduced model is used to estimate noise fluxes, which in turn can be used to estimate a signal-to-noise metric between observed and modelled classes. Classes that exhibit a high signal-to-noise are incorporated into the reduced model.

### Dominance analysis

We modeled allelic dominance as a weighted average of allelic activity:

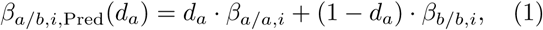

where *β_k/k,i_* refers to the *β* value of the ith isoform in a genotype *k/k*, and *d_a_* is the dominance coefficient for allele *a.*

To find the parameters *d_a_* that maximized the probability of observing the data, we found the parameter, *d_a_*, that maximized the equation:

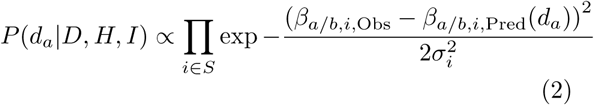

where *β_a/b,i,Obs_* was the coefficient associated with the *i*th isoform in the trans-het *a/b* and *σ_i_* was the standard error of the *i*th isoform in the *trans-*heterozygote samples as output by Kallisto. *S* is the set of isoforms that participate in the regression (see main text). This equation describes a linear regression which was solved numerically.

### Code

Code was written in Jupyter notebooks^31^ using the Python programming language. The Numpy, pandas and scipy libraries were used for computation^32,33,34^ and the matplotlib and seaborn libraries were used for data visualization^35,36^. Enrichment analyses were performed using the WormBase Enrichment Suite^37^. For all enrichment analyses, a *q*-value of less than 1010^−3^ was considered statistically significant. For gene ontology enrichment analysis, terms were considered statistically significant only if they also showed an enrichment fold-change greater than 2.

## Data Availability

Raw and processed reads were deposited in the Gene Expression Omnibus. Scripts for the entire analysis can be found with version control in our Github repository, https://github.com/WormLabCaltech/med-cafe. A user-friendly, commented website containing the complete analyses can be found at https://wormlabcaltech.github.io/med-cafe/. Raw reads and quantified abundances for each sample were deposited at the NCBI Gene Expression Omnibus (GEO) ^38^ under the accession code GSE107523 (https://www.ncbi.nlm.nih.gov/geo/query/acc.cgi?acc=GSE107523).

## Acknowledgements

This work was supported by HHMI with whom PWS was an investigator, by the Millard and Muriel Jacobs Genetics and Genomics Laboratory at California Institute of Technology, and by the NIH grant U41 HG002223. This article would not be possible without help from Dr. Igor Antoshechkin and Dr. Vijaya Kumar who performed the library preparation and sequencing. We would like to thank Carmie Puckett Robinson for the unpublished Dpy transcriptional signature. Han Wang, Hillel Schwartz, Erich Schwarz, Porfirio Quintero and Carmie Puckett Robinson provided valuable input throughout the project.

